# Deep learning based genomic breeding of pest-resistant grapevine

**DOI:** 10.1101/2024.03.16.585323

**Authors:** Yu Gan, Zhenya Liu, Fan Zhang, Qi Xu, Xu Wang, Hui Xue, Xiangnian Su, Wenqi Ma, Qiming Long, Anqi Ma, Guizhou Huang, Wenwen Liu, Xiaodong Xu, Lei Sun, Yingchun Zhang, Yuting Liu, Xinyue Fang, Chaochao Li, Xuanwen Yang, Pengcheng Wei, Xiucai Fan, Chuan Zhang, Pengpai Zhang, Chonghuai Liu, Zhiwu Zhang, Sanwen Huang, Yiwen Wang, Zhongjie Liu, Yongfeng Zhou

## Abstract

Crop pests have profoundly deleterious effects on crop yield and food security. However, conventional pest control depends heavily on the utilization of insecticides, which develops strong pesticide resistance and concerns of food safety. Crop and their wild relatives display diverse levels of pest resistance, indicating the feasibility for breeding of pest-resistant crop varieties. In this study, we integrate deep learning (DL)/machine learning (ML) algorithms, plant phenomics and whole genome sequencing (WGS) data to conduct genomic selection (GS) of pest-resistance in grapevine. We employ deep convolutional neural networks (DCNN) to accurately calculate the severity of damage by pests on grape leaves, which achieves a classification accuracy of 95.3% (Visual Geometry Group 16, VGG16, for binary trait) and a correlation coefficient of 0.94 in regression analysis (DCNN with Pest Damage Score, DCNN-PDS, for continuous trait). We apply DL models to predict and integrate phenotype (both binary and continuous) along with WGS data from 231 grape accessions, conducting Genome-Wide Association Studies (GWAS). This analysis detects a total of 69 QTLs, encompassing 139 candidate genes involved in pathways associated with pest resistance, including jasmonic acid (JA), salicylic acid (SA), ethylene, and other related pathways. Furthermore, through the combination with transcriptome data, we identify specific pest-resistant genes, such as *ACA12* and *CRK3*, which play distinct roles in resisting herbivore attacks. Machine learning-based GS demonstrates a high accuracy (95.7%) and a strong correlation (0.90) in predicting the leaf area damaged by pests as binary and continuous traits in grapevine, respectively. In general, our study highlights the power of DL/ML in plant phenomics and GS, facilitating genomic breeding of pest-resistant grapevine.

## Introduction

The adverse impacts of agricultural pests have long posed challenges for agriculture^1^. Pests, such as aphids, locusts, and moth flies directly damage crops, resulting in reduced yields and compromised quality^2–4^. Other pests like scale insects and thrips indirectly harm crops by vectoring plant viruses^5^. The strong reproductive ability of these pests not only decreases crop production, but also threatens food security^6^. Traditionally, farmers have primarily relied on copious applications of pesticides to control crop pests. However, the excessive use of pesticides contaminated the environment and jeopardized human and animal health^7^. Moreover, the rapid evolution of pesticide resistance among insect populations has progressively diminished the efficacy of these chemicals^8^. Thus, new monitoring methods and control strategies are needed to precisely target the agricultural pests.

Plants and insects have coexisted for over 350 million years^9^. As stationary organisms, plants lack the ability to escape attacks from other organisms, necessitating the utilization of alternative strategies for self-defense. To counter the herbivore attack, plants develop specialized structures or produce secondary metabolites and proteins with toxic, repellent, or antinutritional effects to deter herbivores^10,11^. Plants defend against herbivores both directly, by altering the preferences, survival and reproductive success of herbivores, and indirectly, by emitting volatile organic compounds (VOC) to attract their natural enemies^12^. The defense mechanisms of plants against herbivore attacks involve intricate signal transduction pathways mediated by a network of phytohormones. Plant hormones play a pivotal role in governing plant growth, development, and defense mechanisms^13^. Several plant hormones play roles in both intra- and inter-plant communication in herbivore-damaged plants. The majority of defense responses against insects are triggered by signal transduction pathways involving jasmonic acid (JA), salicylic acid (SA), and ethylene^14–16^. Distinct sets of genes associated with defense are activated by these pathways in response to injury or pest feeding. These hormones can work independently, synergistically, or antagonistically, depending on the specific attacker.

One major pest posing prominent threats to grape yield and quality is the tobacco cutworm (*Spodoptera litura*, Fab, SLF)^17^. In grape cultivation, infestations generally occur between July and September, especially in drought conditions, high humidity, and in proximity to vegetable crops. The larvae of SLF damage the plant mainly by consuming leaf mesophyll tissue, resulting in perforated foliage^18^. Because outbreaks mainly transpire during grape ripening and harvest, extensive feeding can disrupt grape coloring and maturation in the same year. This also impedes nutrient translocation in the following season, leading to poor subsequent bud break and flowering^19,20^. Although a few grape varieties show resistance to SLF, resistant varieties have not been selected and bred to resist SLF infestations. Therefore, determining more resistance germplasm resources, genetics of plant resistance to SLF and the function of resistance genes have been essential for breeding pest-resistant grapevine.

While the throughput and cost-effectiveness of genotyping have improved considerably, the measurement of traits of interest remains insufficient in these regards^21^. Most of the resistance traits have been subjected to visual and sensory evaluations by experienced professional breeders, and phenotypic values of the traits are expressed as qualitative categorical scores^22^. For example, drought stress level (mild, moderate and severe, three categories)^23^ and downy mildew level (seven categories)^24^. An empirical assessment based on the sense of the breeder is not sufficient to evaluate the diverse and continuous variations of the plant resistance. In addition, expertise in visual and sensory evaluation can only be obtained after years of training, and increasing the number of specialized breeders is not practical for large scale and low-cost phenotyping. To further improve the accuracy of GWAS and GS for practical breeding, we need to enhance the data quality of the agronomic traits^25–27^. Deep learning-based image analysis offers a promising approach to address the limitations of current qualitative evaluation methods.

Deep learning is a subset of machine learning that utilizes artificial neural networks to learn from large volumes of data. Deep neural networks contain multiple hidden layers between the input and output layers that enable the model to progressively extract features at multiple levels of abstraction^28^. This hierarchical learning architecture allows deep learning models to learn complex functions and find intricate patterns in data that most traditional machine learning techniques may fail to detect. In agriculture, deep learning has emerged as a promising approach for improving the detection and classification of pests, diseases, and other anomalies. For example, Mohanty et al. developed a deep convolutional neural network able to detect 14 crop diseases and 26 crop species with an accuracy exceeding 95%^29^. Sladojevic et al. used deep neural networks for automated identification of plant diseases from leaf images, achieving an accuracy of 96. 3% compared to 85.9% with a shallow neural network^30^. Overall, deep learning allows predictive models for agricultural applications to achieve greater complexity and abstraction by learning directly from raw data, enabling more accurate and nuanced detection capabilities than traditional analytical techniques. The ongoing explosion of agricultural big data presents an ideal opportunity for deep learning techniques to continue advancing in this area.

In this study, we developed deep learning models that combined object detection and pest severity assessment to generate pest resistance phenotype data from 231 samples of grapevine. Using GWAS, we identified QTL regions associated with resistance and identified pest-resistance related candidate genes within these regions. Furthermore, we integrated GWAS with population genetic analysis, transcriptomic assays, comparisons among different *Vitis* populations, and inter-species comparisons to determine which genomic regions contribute to resistance, as well as to identify potential candidate resistance genes within these regions. Finally, we employed machine learning based GS to evaluate different models in breeding programs of pest-resistance grapevine. Overall, our study contributes valuable insights into the genetics and evolution of pest resistance, which will facilitate genomic breeding of pest-resistant grapevine.

## Results

### Comparing different binary-classification models’ performances through cross-validation

To build the binary classification models for the categorization of damage levels caused by insects, we employed a set of DCNNs, including AlexNet, VGG16, ResNet50, ResNet101, InceptionV3, and DenseNet121. The model performance was rigorously evaluated through cross-validation with the aim of identifying the most optimal model. We measured the complexity of different models using the number of training parameters and assessed their performance, including accuracy and F1 score, through five-fold cross-validation (Supplementary Table 1). The entire cross-validation process utilized a total of 1448 leaf images for training and validation purposes. After training the models for sufficient epochs to ensure convergence, the highest average accuracy was achieved by VGG16 at 90.1%, while the lowest accuracy was observed in ResNet101, scoring 86.9%. Other models fell within the range of 87% to 89% accuracy (Fig. 1a). Correspondingly, VGG16 also exhibited the highest average F1 score, reaching 0.90, while others ranged from 0.86 to 0.89. The strong and significant correlation (*P* = 4.87e-5) between F1 score and accuracy indicates the model’s capability in accurately classifying samples while effectively managing false positives and false negatives (Supplementary Fig. 1). Upon further examination, it was observed that ResNet101, with its deeper network architecture and consequently more training parameters compared to ResNet50, did not exhibit a significant difference in performance between the two models (*P* > 0.05). As the complexity of the model increased, it failed to enhance the performance of this binary classification task. Consequently, we conducted a correlation analysis between accuracy, F1 score, and the number of parameters involved in model training. We discovered a negative correlation trend between the model’s complexity and its performance (Supplementary Fig. 1).

**Fig. 1.**
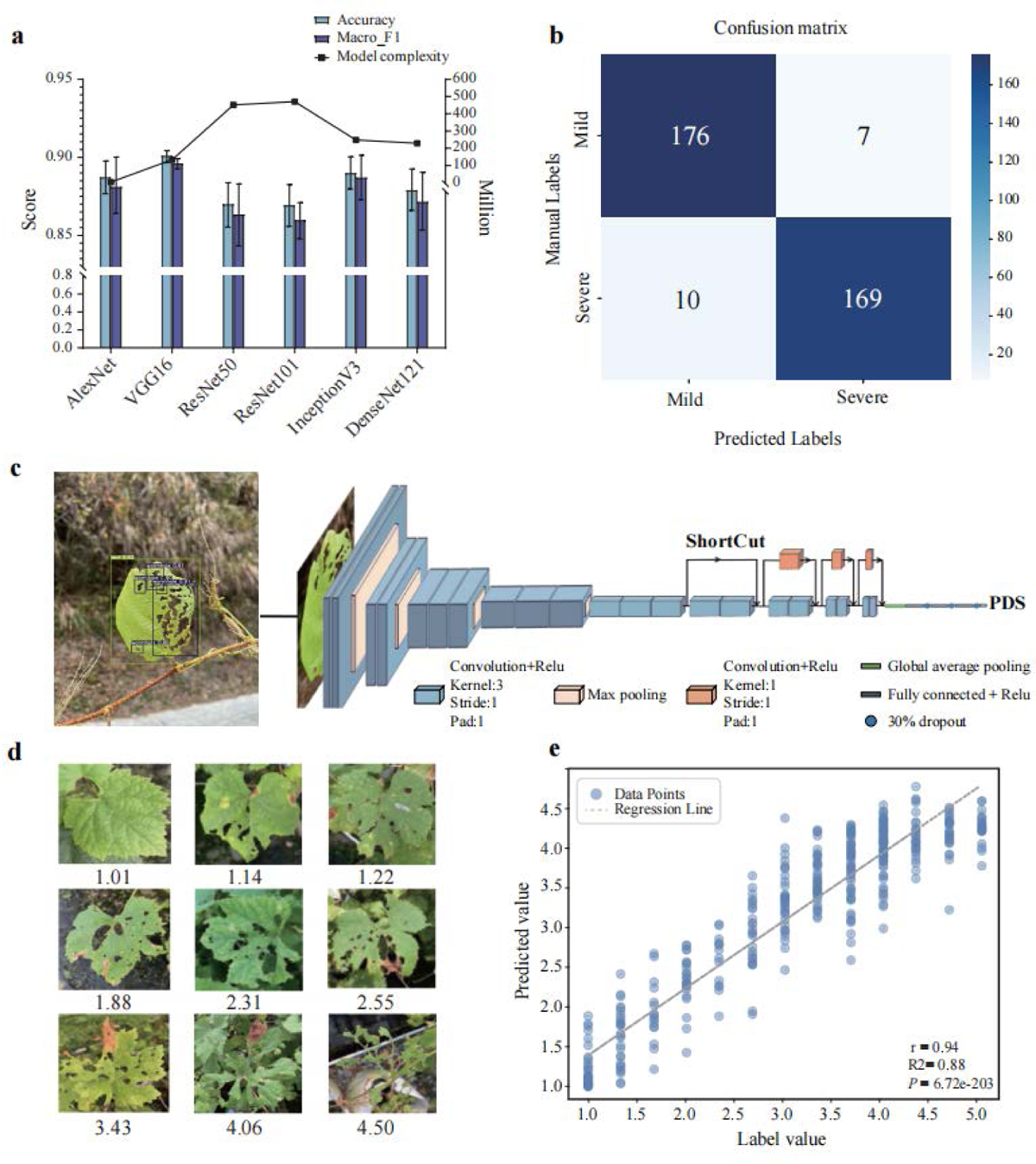
Deep convolutional neural network (DCNN) modeling process and test results. **a**, Performance of six classic convolutional neural networks in cross-validation. The bars correspond to the left Y-axis (accuracy score), while the line is referenced to the right Y-axis (model complexity). **b**, The number of correct recognitions of different categories by VGG16 in the test set. **c**, The model architecture of DCNN-PDS and the workflow for deriving continuous traits from images. **d**, Examples of the DCNN-PDS prediction results for the different extent damages by pest, and the PDS values are shown at the bottom of each plot. **e**, The correlation between the predicted values provided by DCNN-PDS and the manually labeled values in the test set.

VGG16 exhibited the highest accuracy with a relatively low complexity, making it our choice for the binary classification task. Before being deployed, the model’s enhanced performance through transfer learning was assessed on the test set (Supplementary Fig. 2). The VGG16 model achieved an accuracy of 95.3%, precision of 96.0%, recall of 94.4%, and an F1-score of 0.95. The learning curves of VGG16 were shown in Supplementary Fig. 3. Upon scrutinizing the confusion matrix, it was evident that, among a total of 362 test images, the model accurately classified 169 positive samples and 176 negative samples out of 183 (Fig. 1b). It was clear that the VGG16 model adeptly managed the identification of the two categories in the test set. This highlighted the model’s proficiency in generating reliable predictions and indicated its suitability for practical applications in predictive tasks.

### The performance of DCNN-PDS on continuous traits

To further enhance the granularity of damage assessment caused by pests on leaves, we devised and trained a regression model with the ability to assign continuous scores ranging from 0-5 (Fig. 1c). These continuous scores were determined by the extent of pest infestations and reflect the severity of the damage. The pest damage regression model underwent cross-validation and was subsequently assessed using an independent test set. Throughout the cross-validation process, a dataset consisting of 2181 images was employed to assess and determine the best hyperparameter configuration. Utilizing a K-fold cross-validation with K=5, the model achieved an average mean squared error (MSE) loss of 0.1245 and an average mean absolute error (MAE) loss of 0.2343 across various training and validation set distributions (Supplementary Table 2). The fact that the aforementioned metrics were both obtained on the validation set, using the optimal hyperparameters, suggests that the model’s performance remains consistent even when faced with different data partitions, resulting in stable and reliable results (Supplementary Table 2). Throughout the training process, the model was initially set to train for 300 epochs. Because of early stopping, the training was halted after 218 epochs, as the model had reached saturation and continuing training was deemed unnecessary (Supplementary Fig. 4). After DCCN-PDS (pest damage score) converged, the MSE loss on the training set decreased to 0.0241 and 0.0866 for the validation set. The MAE decreased to 0.1061 for the training set and 0.1845 for the validation set, which was far below within one point of the five-point severity scale for pest damage.

Before the model can be applied in practical settings, it was crucial to ensure its strong generalization ability and robustness. To accomplish this, new data specifically for testing purposes was captured, totaling 439 images. These test sets encompassed a variety of grape leaf types and complex backgrounds, introducing variability that differs from the training data (Supplementary Fig. 5). During the testing phase, the model achieved an MSE of 0.1946 and an MAE of 0.3504 on the test set. The DCCN-PDS prediction example showcases the model’s precise quantification of subtle variations in pest damage severity (Fig. 1d). To further analyze the relationship between the predicted values and the manual tag values, a correlation analysis was performed. A significant correlation (*P* < 0.001) between the manual tag values and the predicted values on the test set was observed, as indicated by a Pearson correlation coefficient of 0.94, with a Coefficient of Determination *R*^2^ = 0.88 (Fig. 1e).

### Phenotypic assessment of grape accessions using binary classifications and regression models

To assess the pest-resistance of grapevine populations, we collected cuttings of 231 accessions from National Grape Germplasm at Zhengzhou in Nov. 2021, and cultivated in greenhouse at the Shenzhen station (Fig. 2a and Supplementary Fig. 6). After the outbreak of SLF in the greenhouse in June-August 2022 (Fig. 2a, see Methods), we randomly take pictures for 6-10 leaves for each accession in September 2022 to represent the pest-resistance of the sample. A population structure analysis was performed on a total of 231 grape varieties, including table grape varieties domesticated from *V. vinifera* ssp. *Sylvestris* (EU-Table), table grape varieties originated from hybridization between *V. vinifera* and *V. labrusca* (EA-Table), and wine-making grape varieties domesticated from *V. vinifera* ssp. *sylvestris* (Wine) (Figure 2 and Supplementary Table 3). The examination utilized both the VGG16 model and our custom-developed model DCNN-PDS, resulting in binary and continuous phenotypes (Fig. 2b). Binary classification results revealed 99 varieties exhibited mild damage by pests and 132 varieties have suffered severe pest damage (Supplementary Fig. 7). The EA-table seems to be the least resistant to pests, with 66 out of 92 accessions experiencing severe infestation (50% of the total severe grape accessions). In contrast, the EU-table appears to have stronger pest resistance, with 46 varieties identified as having mild infestation (46.47% of the total instances of mild damage) (Fig. 2d, e). DCCN-PDS predicted and scored damage levels in 0-5 for all accessions based on their images (Supplementary Fig. 7). Regression analysis of the leaf images revealed the EU-table group had the lowest mean and median values at 2.22 and 1.94, respectively (Fig. 2g). Normality testing using the Shapiro-Wilk test indicated the EU-wine group followed a normal distribution (*P* = 0.082), while the *P* values for the other two groups were below 0.05 (Supplementary Fig. 8). Homogeneity of variance among the three groups was supported by a Bartlett test (*P* = 0.5542). Given these results, the Wilcoxon rank sum test was used to detect differences between groups and found significant differences between all three. It was evident that, in comparison to EA-table and EA-wine, EU-table displayed significant (*P* < 0.001) phenotypic distinctions, signaling a generally heightened pest resistance when contrasted with the aforementioned two populations (Fig. 2g). Phylogenetic regression was performed by mapping regression phenotypes onto the phylogenetic tree (Fig. 2b), and correlation and Spearman^’^s footrule distance were calculated. The correlation between regression value order and tree order was 0.2747, with a Spearman^’^s footrule distance of 19405. In contrast, the average correlation of random sequences was −3.09e-4, with an average distance of 23102. Single sample t-tests showed both analyses were significantly non-random (*P* < 2.2e-16), suggesting a significant association between pest resistance and phylogenetic evolution.

**Fig. 2.**
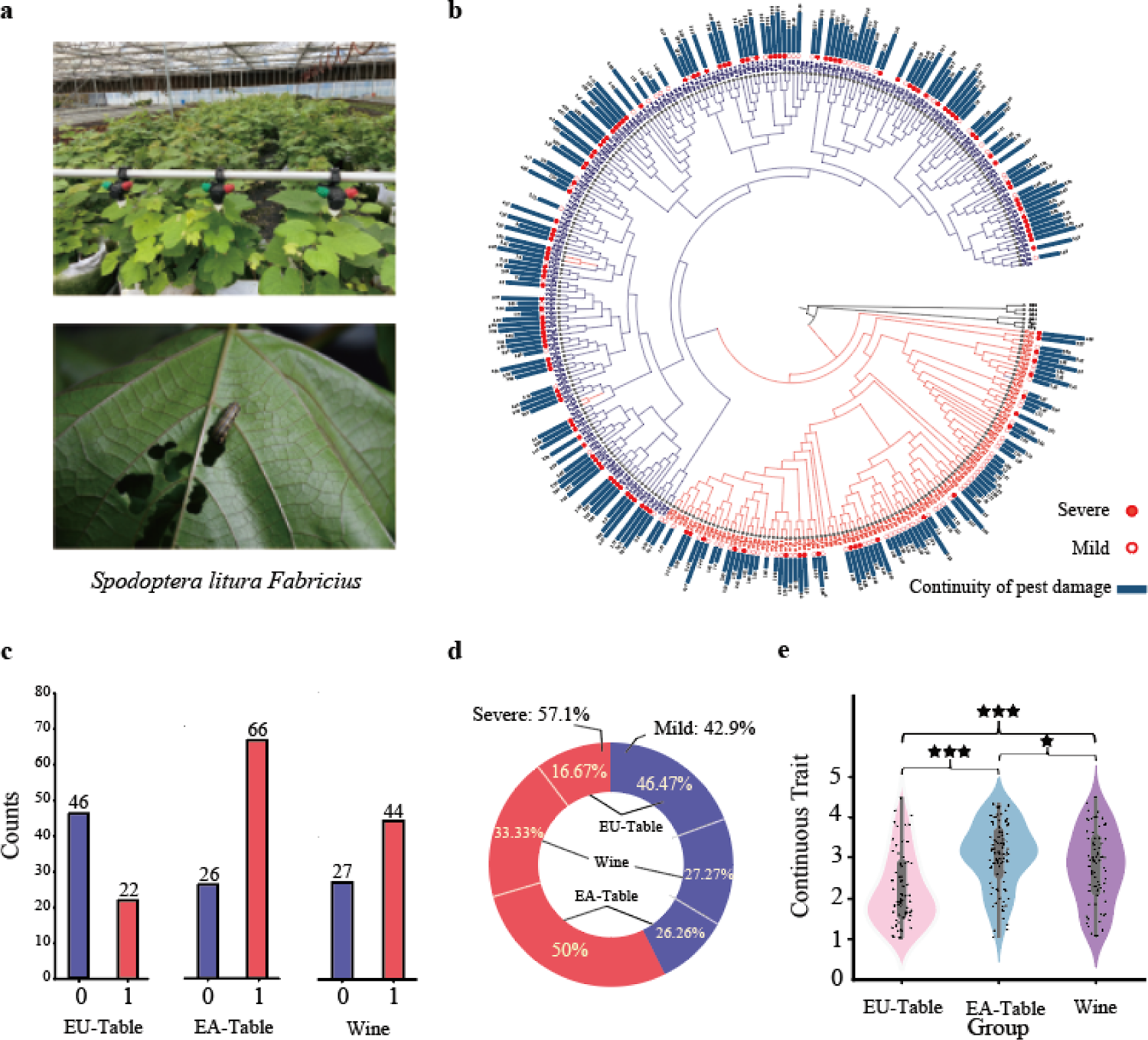
Statistical analyses the overall phenotype of grapevine accessions. **a**, The greenhouse conditions and the experimental setup for SLF infestation. **b**, The evolutionary tree of 323 individuals is divided into two populations: Europe and America, and Eurasia. The bar chart displays continuous trait values for each accession, with solid red dots indicating severity in binary traits. **c**, Three categories have been classified based on the purpose of grapes, each corresponding to the distribution of binary phenotypes. **d**, A pie chart depicting the overall binary classification phenotypes, and described the percentage of their respective binary phenotypes. **e**, Violin plots depicting the distribution of continuous phenotypes for the three categories, with significance differences denoted by asterisk (★: *P*<0.05, ★★★: *P*<0.001).

### Genome-wide Association Study (GWAS) of binary and continuous pest-resistant traits

To further map the quantitative genetic basis of pest damage, we conducted GWAS for binary traits derived from VGG16 and for continuous traits obtained from DCNN-PDS, respectively (Fig. 3a and Supplementary Fig. 9). This aimed to investigate their ability to efficiently identify genetic loci associated with pest resistance. The analysis revealed 33 significant QTLs associated with binary traits, involving 67 candidate genes, and 36 QTLs associated with continuous traits, encompassing 85 candidate genes (Supplementary Tables 4 and 5). Genes associated with continuous traits, as identified through GO enrichment analysis, were found to be primarily involved in the calcium ion binding process (*P* < 0.05). In contrast, the genes associated with binary traits did not exhibit significant enrichment in molecular functions, nor did the total 139 non-redundant candidate gene sets (Supplementary Table 6). Notably, all three gene sets detected a cluster related to involvement in protein kinase activity and protein phosphorylation, although the significance level was not sufficient (Supplementary Table 7). We found the *1-aminocyclopropane-1-carboxylate oxidase homolog 1* (*ACO*, *Vitvi008234*) and *sterol 3-beta-glucosyltransferase UGT80B1* (*TT15*, *Vitvi022534*) located on chromosome 5 and 12 of binary phenotype results, and *probable protein phosphatase 2c 14* (*PP2C*, *Vitvi000748*) located on chromosome 1 of continuous phenotype results which has shown related to pest resistance in plant in previous research (Fig. 3a). *Vitvi008234* and *Vitvi022534* are involved in the synthesis of ethylene and flavonoids (Supplementary Table 8), respectively^31,32^. These two compounds play a significant role in influencing the plant’s defense against herbivore feeding^33,34^. *Vitvi000748* encodes a PP2C-type phosphatase (Supplementary Table 9), PP2C has been identified as a negative regulator of protein kinase cascades activated in response to stress^35^, existing research has indicated that PP2C is involved in the plant’s defense response against herbivores^36^. In addition, we identified a strong signal (Chr16: 26.7M-27.0M) for continuous trait (snp_16 26926513, *P* =1.71e-06). Plants possessing both pure and mutated loci demonstrate lower scores and diminished pest damage (Fig. 3b). The predominant locus among the three populations is characterized by heterozygous haplotypes, with a higher prevalence of homozygous mutant haplotypes in the EU-Table and an increased frequency of homozygous mutant haplotypes in the EA-Table (Fig. 3c). Additionally, the genome-wide analysis of genetic differentiation (*F*_ST_) and genetic diversity (π) indicates an absence of selection events in this region (Fig. 3d). The linkage disequilibrium analysis highlights a significant linkage in the promoter region of *CDPK-RELATED KINASE 3* (*CRK3*, *Vitvi031045*) (Fig. 3e). The gene *Vitvi031045* encodes a Ca^2+^/CaM-dependent protein kinase related protein kinase and participate in CaM-mediated signaling pathways^37^. (Supplementary Table 7). Previous studies have demonstrated that CRK overexpression and knockout mutants exhibits increased and decreased defense traits against SLF^38^. Furthermore, it was found that the transcription of *CRK3* is positively regulated by JA and abscisic acid (ABA) signals, particularly under herbivory-induced stimulation^38^. It plays a pivotal role in modulating plant defense responses by phosphorylating and activating *WRKY14*, and enhancing the defense response through the upregulation of *PDF1.2* ^38^.

**Fig. 3.**
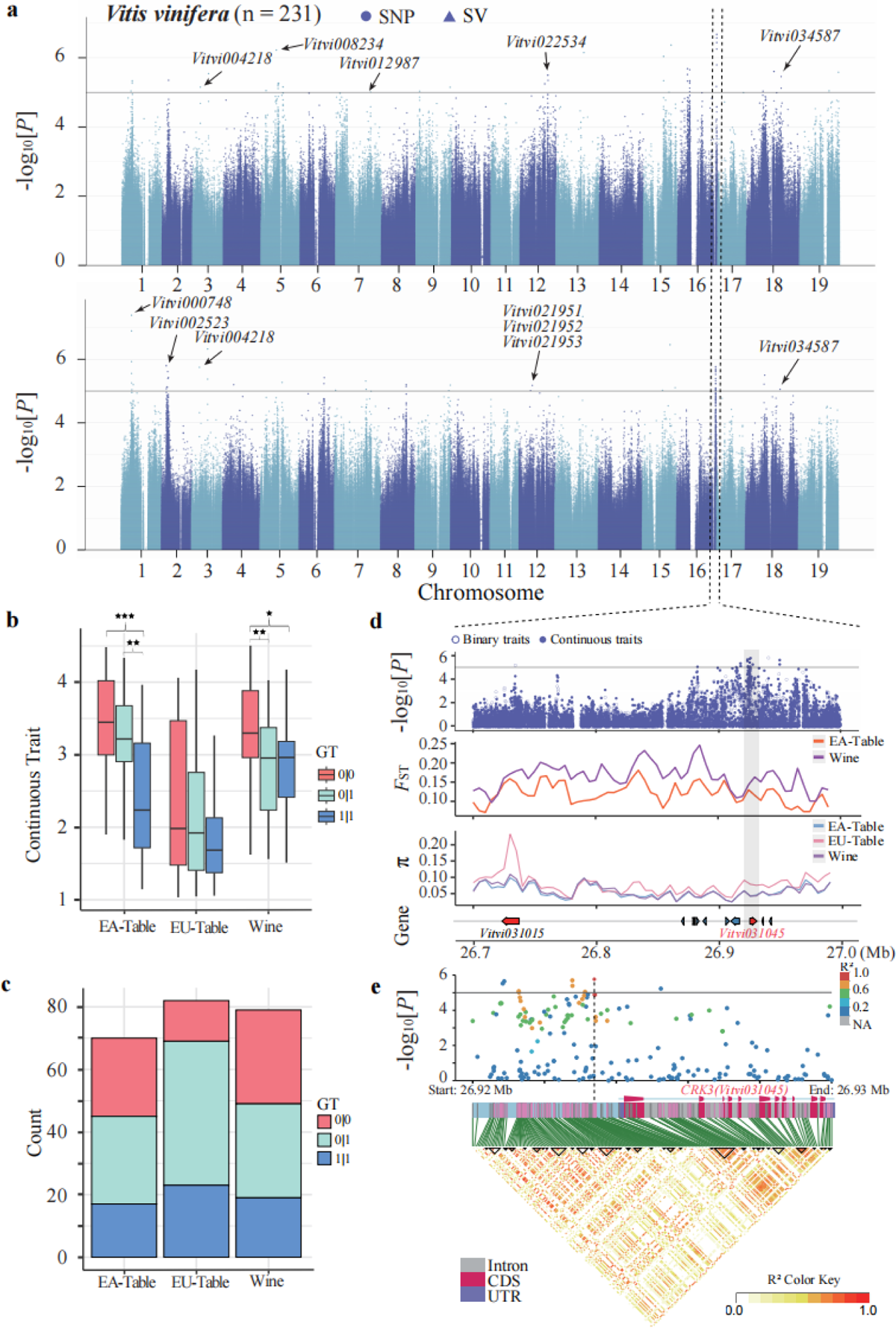
GWAS results based on the phenotypes generated by the DL model. **a**, The Manhattan plot reveals GWAS results for two phenotypes, with the upper section representing binary phenotype results and the lower section representing continuous phenotype results. **b**, A boxplot depicting the continuous phenotype corresponding to the genotype of snp16 26926513 in three grape populations, asterisk indicate significant differences between them (★: *P*<0.05, ★★: *P*<0.01, ★★★: *P*<0.001). **c**, The number of individuals corresponding to three genotypes in three populations, where 0/0 represents no mutation, 0/1 represents heterozygous mutation, and 1/1 represents homozygous mutation. **d**, The region on chromosome 16 (chr16: 26.7-27.0 Mb) was zoomed in, sequentially illustrating the corresponding GWAS results, *F*_ST_, π, and the situation of the corresponding genes. **e**, Linkage disequilibrium analysis was conducted for the gene *Vitvi031045* and an upstream segment (chr16:26.92-26.93 Mb), with the upper section revealing the *P* values of the GWAS results.

### Integration of GWAS and RNA-seq identified pest-resistant candidate genes

In order to verify the accuracy of the DL model predictions and GWAS analysis results, we conducted transcriptomic analyses. Under assault by the two-spotted spider mite, *Tetranychus urticae*, another herbivorous insect, we identified a total of 3979 differentially expressed genes (DEGs) with *P* <= 0.05 and a fold change greater than 2. Subsequently, a total of 2010 upregulated genes and 1969 downregulated genes were identified (Fig. 4a and Supplementary Table 10). GO enrichment analysis revealed these genes are significantly involved in signal transduction, defense responses, and metabolic processes, consistent with previous studies^39^ (Supplementary Table 11). It is noteworthy that DEGs were also found to be enriched in a cluster with the highest significance, specifically in protein phosphorylation and protein kinase activity (protein serine/threonine/tyrosine kinase activity) (*P* < 0.05), in line with our GWAS analyses. We observed a convergence between the DEGs uncovered through RNA-seq and the genes associated with binary traits (BTGs) and continuous traits (CTGs). There were 13 genes common to both traits, with 7 genes shared between BTGs and DEGs, and 12 genes shared between CTGs and DEGs, respectively (Fig. 4b). Notably, all three analyses identified the presence of two genes (*Vitvi034587* and *Vitvi031046*). *Vitvi031046 (SAUR32)* encode an auxin-responsive protein and *Vitvi034587 (ERF110),* known as *ethylene-responsive transcription factor* (*ERF110, Vitvi034587*), which was significantly induced under conditions of herbivore attack (Fig. 4c and Supplementary Table 12). *Vitvi012987* (*ATHB-6*) and *Vitvi031015* (*DMR6*) constituted the intersection of BTGs and DEGs, playing roles in the regulation of the ABA signaling pathway and the synthesis of defensive oxygenase, respectively^40,41^. They were also significantly induced in expression, indicating their relevance to pest defense (Fig. 4c). Within the intersection of CTGs and DEGs, two key genes have been identified (*Vitvi021951* and *Vitvi021953*). These genes are associated with continuous traits (snp_12 9025204) in GWAS and are strongly induced under herbivore attack. Additionally, it is noteworthy that these three genes are tandemly linked (Chr12: 9.01M-9.04M) and are encoded by the same gene, *calcium-transporting ATPase 12, ACA12* (Fig. 4d). *ACA12* encodes a calcium ion pump localized on the plasma membrane of plant cells (Fig. 4e). Mutants of *ACA12* have been demonstrated to exhibit extreme sensitivity to herbivorous attack by *Spodoptera littoralis*. This sensitivity manifests as a significant increase in leaf damage, accompanied by a decrease in intracellular calcium ion concentrations^42^. Simultaneously, *Vitvi031045* (previously mentioned in the GWAS results) was also found to be significantly induced (Fig. 4c, e). Analysis of integrated transcriptome data demonstrated the effectiveness of the identified pest resistance genes across both traits through the application of DL models.

**Fig. 4.**
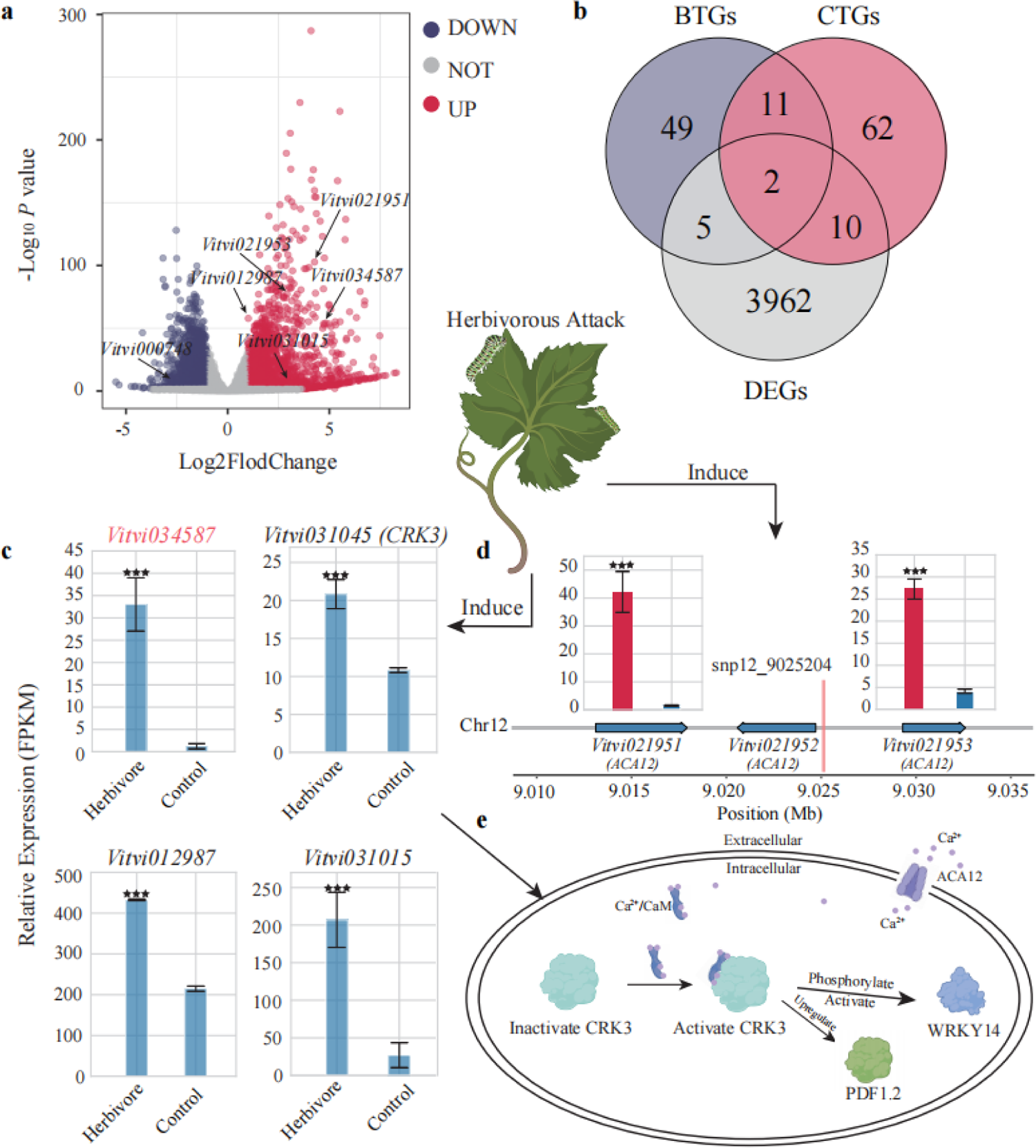
Differential gene expression under herbivorous conditions in grapevine. **a**, Volcano plot of gene expression differences, with red indicating upregulation and blue indicating downregulation for each gene, the genes indicated by the arrows illustrate that they are also located by the GWAS results. **b**, Venn diagram illustrating the overlap between genes associated with binary and continuous phenotypes, as well as differentially expressed genes. The numbers represent the count of genes. **c**, Gene expression patterns of the four pest-defense genes which derived from the intersection of GWAS and RNA-seq results. **d**, Three genes in the region of chromosome 12 (9.010M-9.035 Mb) encode the same protein and were detected by GWAS (snp12_9025204). Bar charts for each gene depict the corresponding expression levels, where red indicates herbivory conditions, and asterisk denote significant differences (★: *P*<0.05, ★★★: *P*<0.001). **e**, The functions performed by two insect-resistant genes within the cell are elucidated, along with a description of a potential interaction between them.

### Improving genomic selection with the optimal machine learning model

The outcomes from GWAS with binary and continuous traits were utilized to construct genomic selection models based on machine learning methodologies, respectively. Among four evaluated models, including Logistic Regression, Support Vector Classifier (SVC), Random Forest Classifier (RFC) and Naïve Bayes Classifier models for the binary trait prediction (Supplementary Table 13). The performance of the RFC model with different variations consistently fell short, achieving an accuracy that did not exceed 85%. Naive Bayes outperformed RFC but fell behind the other two models, achieving a maximum average accuracy of 87.6%. As the number of variations increased, the performance of the SVC model consistently improved. Beyond 500 variations, the average accuracy showed a gradual plateauing, reaching its peak at 93% when the number of variations reached 5000. The logistic regression model was deemed more suitable for this modeling task. In contrast to the SVC model, logistic regression reached its peak accuracy at 2000 variations (94.5%) and exhibited a declining trend in accuracy with 5000 variations (Fig. 5a). We performed the final validation of the test set by combining the logistic regression model with the appropriate number of variations. The accuracy achieved was 95.7%, and a significant correlation of 0.94 (*P* = 1.4e-11) was observed between the predicted values and the actual phenotypes (Fig. 5b). In the prediction of continuous traits, we compared various linear models (Lasso, Ridge, ElasticNet) and machine learning models, including Random Forest Regression (RFR) and Support Vector Regression (SVR) (Supplementary Table 14). Among these models, the Lasso model exhibited a poor performance, with the highest average correlation coefficient not exceeding 0.72. The accuracy of the RFR showed negligible variation, reaching a maximum of only 0.79. ElasticNet performed well, achieving a high accuracy of 0.88, and demonstrated stable performance beyond 500 variations. According to the results, the Ridge model exhibited instability, with a sudden increase in accuracy after finding an appropriate number of variations, reaching a peak of 0.89 at 10,000 variations. SVR outperformed all other models, showing a steady increase in performance. Beyond 2000 variations, its performance gradually stabilized, reaching a peak accuracy of 0.92. (Fig. 5c). Based on the results of cross-validation, we selected SVR and applied the identified 10,000 SNPs to predict the continuous phenotype in the test set. Ultimately, we obtained a significant correlation coefficient of 0.90 (*P* = 3.5e-9) between the predicted and actual phenotypes, demonstrating the accuracy of the predictions (Fig. 5d). Additionally, all Support Vector Machine models (SVC, SVR) undergo cross-validation to select the optimal kernel function that represents the corresponding optimal results (Supplementary Fig. 10).

**Fig. 5.**
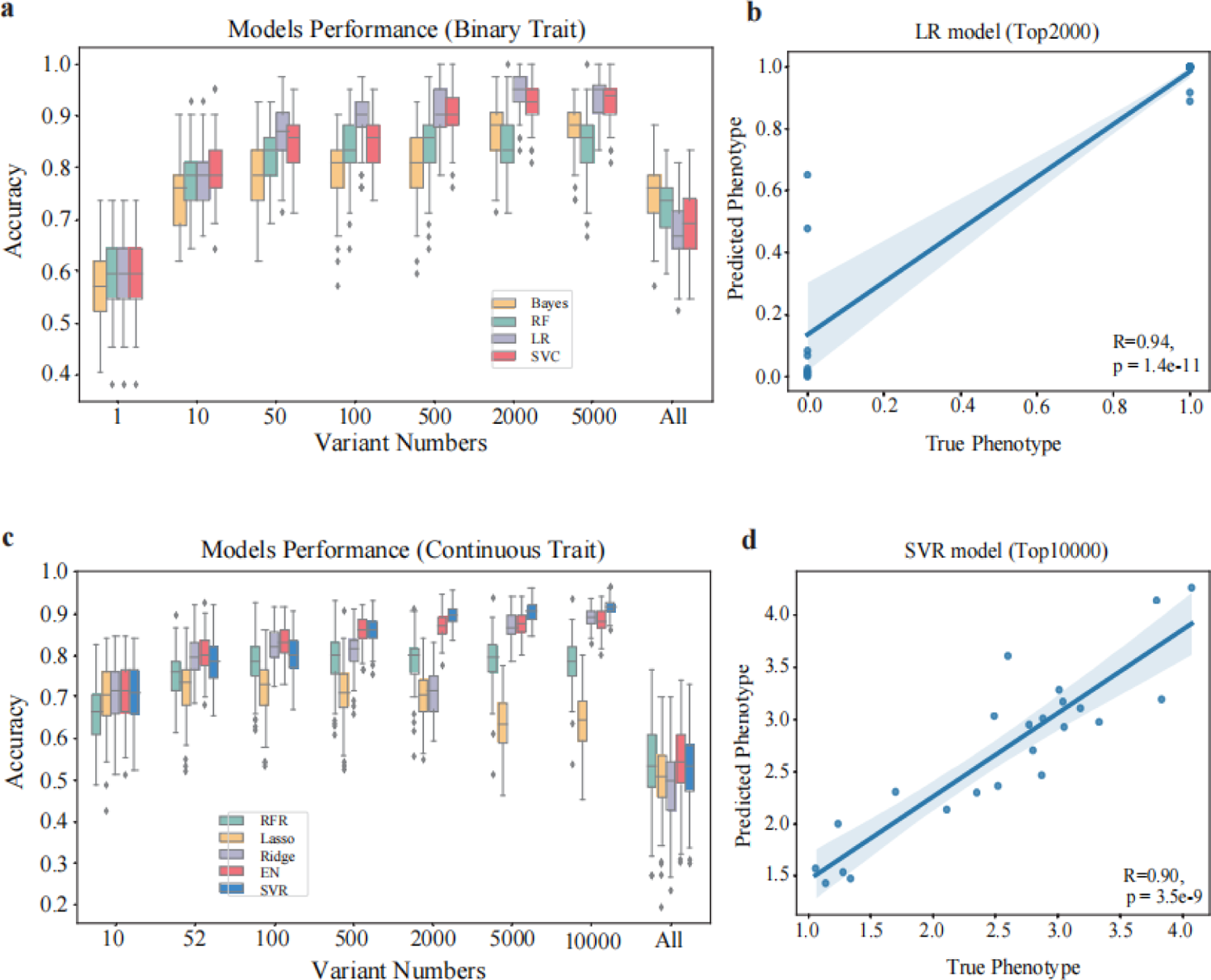
Machine learning based genome selection of pest-resistance in grapevine. **a**, Cross validation of the accuracy of the four classification models, using the average accuracy value as the accuracy indicator. **b**, Using the optimized model and variable count, predictions were made on the test set samples, revealing the correlation between the predicted binary phenotype and the true binary phenotype. **c**, Cross validation of the accuracy of the five regression models, using the average correlation coefficient value as the accuracy indicator. **d**, Using the optimized model and variable count, predictions were made on the test set samples, revealing the correlation between the predicted continuous phenotype and the true continuous phenotype.

## Discussion

Our study implemented cutting-edge breeding methodologies to develop grapevine cultivars with enhanced resistance to pests, leveraging recent breakthroughs in DL based plant phenomics and ML based genomic selection. Specifically, we employed GWAS to map the genetic basis of pest resistance in grapes^43,44^. Subsequently, genomic selection is utilized to selectively breed individuals harboring pest-resistant alleles. Although binary phenotypes are commonly deployed in pest resistance analyses, they overlook subtle-effect genes in GWAS. Conversely, the continuous phenotype generated by DCCN-PDS enables the detection of minor-effect genes, thereby offering a more comprehensive understanding of pest resistance mechanisms. We integrated DL based binary and continuous pest damage traits, leading to the identification of 139 candidate genes endowing pest resistance, facilitating machine learning based genomic prediction of pest damage traits in grapevine breeding.

### The genetic basis of pest-resistance in grapevine

This study aimed to identify genes associated with resistance to *Spodoptera litura Fabricius* (SLF) in grape. Since there was no prior knowledge about the impact of identified genes on SLF resistance, both binary and continuous damage scores were used as phenotypes in GWAS. Our findings revealed 139 candidate genes related to insect resistance that participate in various signaling pathways, including jasmonic acid (JA), salicylic acid (SA), ethylene, and others. Notably, the gene *CRK3* showed a significant association with both types of phenotypes, indicating its crucial role in SLF resistance. Previous research has also suggested *CRK3*’s involvement in plant defense responses, as overexpressing *CRK3* in Arabidopsis enhances defense properties, while knockout mutants exhibited reduced expression levels of the defense gene *PDF1.2*^38^. Additionally, *CRK3* has been demonstrated to phosphorylate tyrosine residues of herbivory-responsive regulators like *WRKY14*, highlighting its participation in the defense response against SLF^45–47^. It is worth mentioning that *ACA12*, another gene associated with continuous damage scores, encodes a calcium ion pump protein localized to the plasma membrane of plant cells^48^. Research has shown that *ACA12* mutants are more susceptible to SLF attacks compared to wild-type plants, suggesting that *ACA12* may play a vital role in providing a calcium ion-enriched environment for *CRK3*^42^. This interaction could potentially activate *CRK3* or other defense proteins reliant on calcium ion signaling, initiating subsequent defense responses^38^. Nonetheless, further investigation is necessary to validate this hypothesis.

It is interesting to note that genes conferring resistance to diseases tend to display higher levels of polymorphism compare to other genomic regions, likely due to the selective pressure imposed by rapidly evolving pathogen populations^48^. Unfortunately, only a minuscule proportion (approximately 0.1%) of the available biodiversity in resistance loci controlling pests and pathogens has been leveraged in commercial crop varieties. To develop more pest-resistant crops, integrating diverse genetic resources from local varieties, wild ancestors, and wild relatives into breeding programs is essential^49^. These resources possess distinct combinations of resistance traits shaped by evolutionary pressures, rendering them valuable sources of novel genetic materia^50,51^. By capitalizing on the innate resistance of host plants through these resources, breed selection becomes more effective, and germplasm utilization is significantly improved, ultimately facilitating crop improvement.

### DL/ML based methods for grapevine genomic breeding

Compared to traditional methods, deep learning enables more efficient, objective and accurate assessment of phenotypic variation in plants^52,53^. Commonly, the extent of leaf damage is manually assessed by visual inspection therefore it is slow, labor-consuming and subjective to the judgement of individual assessors. Our DL models can objectively assess one leaf image within 0.03 to 0.04 seconds and measure its damage condition with high accuracy, up to 0.95 for binary phenotype and with a high correlation, up to 0.94 for continuous scores, compared to the manual labeling. Therefore, the sample size can be easily scaled up if required. Our DL models improve the feasibility to establish large scale phenomics and empower the molecular breeding of grapes^21^. Noteworthily, in insect pest research, DL is mostly applied to identify the pest types based on the morphological features of the pest itself^54^, hence not designed to evaluate the pest resistance of the plants. To precisely determine the resistance feature of plants themselves, it is required to measure the damage conditions in various infestation parts of the different plant samples, for example damage to air potato tubers^55^, leaf damage^56^, dead heart tillers and whitehead-panicles^57^, roots damage^58^ and so on. In our study, the major pest in grape cultivation is SLF. Its larvae mainly consume grape leaves and can cause complete defoliation^59^. Therefore, our analysis focused on damages to the grape leaves, which negatively corelated with SLF resistance for different grape varieties. Generally, the damages of plants caused by the disease and pest are often considered together in most studies using popular DL models designed for image recognition^54,60,61^. It is difficult to disengage the disease and pest resistant capabilities from the same plant. To avoid this, our study is deliberately based on disease-free plants and our model is then capable to measure the damage (mild vs. severe) caused by the pest only. We selected VGG16, which outperformed the other widely used image recognition models, including AlexNet, ResNet50, ResNet101, InceptionV3, and DenseNet121, as our DL models backbone. We added four residual networks based on the backbone to generate both binary and continuous damage scores. For genomic selection of pest-resistant grapevines, we compared multiple ML methods including random forest, naive bayes, support vector machines, elastic net and other models, to avoid empirical bias in model selection. We finally selected logistic regression (95.7 % in accuracy) and support vector regression (0.90 in correlation coefficient) as models for precisely predicting binary and continuous phenotypes, respectively.

In conclusion, our results indicate that deep learning is an effective technique for phenotypic evaluation with high speed and low biases. It lays the foundation for large-scale phenomics data establishment. Combined with the well selected ML model with high prediction accuracy, our study is able to uncover valuable alleles that enhance pest resistance and facilitating efficient breeding for pest-resistant grapevines. Our integrative analysis of genomics, phenomics, transcriptomics showed the power of genetic decode of grape pest damage. It empowers the genomic breeding of grape, which greatly reduce the breeding time per generation. The phenomics evaluate model and genomic selection model would be integrated to automated breeding platform. After that, the breeding strategy would be driven by mass breeding data and big data technology. Artificial intelligence (AI) breeding will be the next step for grape breeding and renovation the breeding strategy.

## Materials and Methods

### Viticulture conditions and pest infestations outbreak

A total of 231 grape varieties were introduced from the Grape Germplasm Repository of the Zhengzhou Fruit Research Institute of the Chinese Academy of Agricultural Sciences and planted in the Grape Germplasm Resource Nursery of the Shenzhen Institute of Agricultural Genomics (AGIS) of the Chinese Academy of Agricultural Sciences (CAAS) in Nov. 2021. All experimental varieties were 2-year-old cuttings from standard vines and were cultivated in pots under greenhouse conditions, with each pot including at least three branches to represent given accession. All plants were subject to standard management practices prior to the experiment, including cultivation, irrigation, fertilization, pruning, and disease control (Supplementary Fig. 6). To examine the impact of different grape varieties on pest damage, we randomly arranged different grape varieties in a greenhouse. We performed a ∼20 days experiment, when a natural outbreak of tobacco cutworm occurred in our greenhouse during July-August 2022, with 9-10 larvae per plant. The pesticide was sprayed after the experiment ended (Fig. 2). We randomly take pictures for six to ten leaves for each accession in early September 2022 to represent the pest damage of the sample.

### Collection and preprocessing of image datasets for model training

The images were captured using the high-quality built-in camera of a Sonya ZV-1 II. The camera incorporates state-of-the-art image sensor technology and an image processing chip, ensuring exceptional image quality and precise detail capturing. This approach was not restricted to specific grape varieties or background environments. Instead, the training images were randomly captured, encompassing a variety of scenarios and conditions to better adapt to the complexity encountered during prediction. We captured thousands of grape leaf samples from various locations, including the wilderness and farms (https://github.com/zhouyflab/Pest-Resistance). These samples exhibited diverse colors, backgrounds, and varying degrees of pest damage. To locate the main portion of the leaves and reduce unnecessary pixel information, we trained an additional YOLO object detection model specifically for leaf detection^62^, which enabled us to accurately and automatically truncate the target (leaf) bounding boxes from thousands of images (Fig. 1c). Although the model has the capability to detect wormholes as well, they were not considered in this study, and the weights file for this YOLO model can be available for download (https://github.com/zhouyflab/Pest-Resistance). The object detection model achieved an impressive mAP value of 0.91 after undergoing 300 epochs of training (Supplementary Fig. 11). The leaf images were loaded and resized to a fixed size of 224×224 pixels using the Python Imaging Library (PIL) (Version:9.4. 0) to ensured that all input images had a consistent size for subsequent processing and model input, then normalize the images by scaling the pixel values to the range of 0-1. To increase the diversity and generalization ability of the training data, data generators were created using the ImageDataGenerator class from Keras library (Version:2.4. 0) which performed random transformations and augmentations on the training data, including random rotation within a range of 45 degrees, random width and height shifts of 15%, horizontal flipping, and random zooming with a range of 50%^63^.

### Image labeling and dataset splitting process

To train the binary classification model, a total of 1810 images were labeled by using one-hot encoding, the leaves in the image with pest damage exceeding 25% were labeled as 1, while leaves with damage below or equal to 25% were labeled as 0, indicating mild damage. The dataset was randomly split into a 4:1 ratio, with 80% of the data used for cross-validation (training and validation), and the remaining 20% used as the test set. After selecting the most optimal model, we further divided a subset of 100 images for validation, while the remaining images were designated as the training set. This division was implemented to enhance the model’s effectiveness during the formal training process. For the regression model, a total of 2620 images were utilized for training and evaluation. Among them, 439 images were set aside as a test dataset, while the remaining 2181 images were involved in the training and cross-validation process, following a 4:1 ratio splitting. All the images are assigned label values ranging from 1 to 5. When the leaf is damaged by pests within 20%, it is assigned a label value of 1, for every subsequent 20% increment in damage, the label value is increased by one, the highest label value of 5 corresponds to damage exceeding 80%. Note that all labels, including binary classification labels and regression labels, are determined among three raters to reduce subjective bias (Supplementary Fig. 12).

### Modeling process of six binary task models

All of the deep learning model that recognize grape leaf is built that based on the Keras library with TensorFlow (Version: 2.4. 0) backend^64^. All modeling procedures were conducted on a Linux platform with a GeForce RTX 3090 GPU, equipped with 24 GB of memory.

For the binary-classification modeling procedures, a comparative analysis of the performance of the AlexNet^65^, VGG16^66^, ResNet50, ResNet101^67^, InceptionV3^68^, and DenseNet121^69^ models was conducted for the classification of pest damage into two categories: mild and severe. The network architecture of these models was modified with a specific emphasis on the fully connected layer and the output layer. In the fully connected layer, all models were composed of two fully connected layers with relu activation function, each consisting of 4096 neurons. Additionally, a dropout rate of 0.5 was applied to enhance the model’s robustness^70^. The activation function used in the output layer was sigmoid, which was chosen to align with the binary cross-entropy loss function. To prevent overfitting, early stopping was implemented to terminate the training process when it no longer resulted in meaningful improvements. The accuracy on the validation set was monitored as the indicator for early stopping. The models exhibited varying levels of complexity, leading to different patience values for performance improvement. For example, AlexNet was tolerated for 20 epochs, while DenseNet121 required 40 epochs. The batch size was set to 64, and each batch was generated by the data generator with real-time data augmentation^63^. To evaluate these models, we employed K-fold cross-validation with K=5 and used the scikit-learn library (Version: 1.0.2) to calculate accuracy and F1-score metrics as evaluation criteria. During cross-validation, the evaluation indicator for each model is its average performance metric, which helps reduce the impact of randomness introduced by a single partitioning of the dataset. Random seeds were applied to ensure consistent data splitting during the cross-validation process across all models. All the six models share the same hyperparameters, including a learning rate of 1e-4 and the utilization of the Adam optimizer^71,72^ for 200 epochs of training. The entire cross-validation training process starts from scratch. Once the optimal model is selected, the hyperparameters remain unchanged, and transfer learning is applied to enhance model performance^73–75^. Subsequently, the model is tested on the test set.

### DCNN-PDS modeling procedures

As for the regression model, the model gives a continuous value to evaluate the pest-damage score (PDS) of grape leaves. The modeling process of this model goes through Keras library^64^ (Version:2.4. 0) and is completed on the Linux platform. The network architecture is constructed based on a combination of VGG16^66^ and residual networks^67^. VGG16 is known for its excellent feature extraction capabilities. However, simply stacking networks to enhance model performance may result in the loss of important image information and degrade the overall network model^67^. To address this concern, we integrated the concept of residual networks. By incorporating residual blocks, it becomes possible to stack deeper convolutional neural networks (CNN) without excessive concerns about network degradation or related issues. Four residual blocks were introduced after the convolutional layer of VGG16, each residual block consists of two convolutional layers with 512, 256, 128, and 64 filters respectively, and then a global average pooling (GAP) layer was added behind the last convolutional layer which help to reduce the feature dimensions. Three fully connected (FC) layers with relu activations were appended behind the GAP layer to do the feature extraction and fusion. In more detail, each of the three FC layers had 1024 neurons. After each FC layer, a dropout layer with a dropout rate of 0.3 was added to reduce overfitting. The network architecture deviated from the original VGG16, which was designed for multiclass classification with softmax activation in the last FC layer^66^. Instead, the objective was to address the numerical regression problem of pest-damage scores. Therefore, sigmoid activation was used in the last FC regression layer^76^, then finally, the output value of the last FC layer X5 was used as the pest impact score which is corresponded to the value of the manual label. (Fig. 1c)

It is worth noting that the modeling process of the model utilizes transfer learning^73^. Compared to training the model from scratch, the pretrained model has already been trained on large-scale datasets, enabling it to possess more powerful feature extraction and generalization capabilities^74^. In this study, the parameters of the convolutional layers of DCNN-PDS have been pre-trained on ImageNet^75^ and are frozen during the training process^77^. For this regression model, the mean squared error (MSE) was chosen as the loss function to be optimized during the training process^76^. Additionally, the mean absolute error (MAE) was used as a performance metric to evaluate the model’s accuracy. By employing K-fold cross-validation K=5, the model utilizes the Adam optimizer^71^ with a learning rate of 1e-3. The batch size was set to 64, the overall training process consisted of 300 epochs, and an early stopping mechanism (patience = 35) was additionally implemented to avoid unnecessary training. After completing the cross-validation evaluations, we proceeded with the formal training of the model and perform final evaluation of the model on a brand-new test set while maintaining optimal hyperparameters unchanged. To further assess the performance of the model, we utilized the Numpy library (Version:1.19. 5) to calculate the Pearson correlation coefficient^78^ between the predicted values and the labels, aiming to determine their linear relationship.

### Grape phenotype calculation

Six to ten images are available for each grape accession to determine its phenotype (https://github.com/zhouyflab/Pest-Resistance). To calculate the binary phenotype, indicating the severity or mild of pest damage, the model computes a probability value of the 0-1 classification for each image. Images with an average probability value below 0.5 are classified as mild pest damage, while those with an average probability value greater than 0.5 are classified as severe pest damage. For a more detailed evaluation of pest damage, the DCCN-PDS provides direct damage scores for multiple images. These scores are then averaged to obtain the final continuity phenotype value. The accurate scores provided by DCCN-PDS quantify the severity of pest damage on a continuous scale from 0-5, without dimensions. This approach offers a more objective alternative to manual measurement and provides data suitable for genetic analysis.

### Distribution statistics of pest phenotypes

The Sapiro-Wilk test was employed to assess the normality of the sample data. Specifically, the shapiro. test was utilized in R to compute the W-statistic and associated p-value for various samples. A p-value inferior to 0.05 signifies that the sample’s phenotypic distribution conforms to a normal distribution. In order to evaluate variance homogeneity on normally distributed data, the Bartlett test was performed. The bartlett. The test function in R was leveraged to determine the corresponding p-value for each sample. If the p-value surpasses 0.05, it may be inferred that the variance of the data exhibits homogeneity. We employed R’s Spearman and rcorr functions to calculate Spearman’s footrule distance and correlation, respectively. List A was based on the order of the phylogenetic tree, while List B was based on the degree of pest damage to different varieties. Furthermore, we generated 100 random orders for conducting a single-sample t-test.

### The GWAS analyses

In this study, paired-end resequencing reads were mapped to the V. vinifera reference genome PNT2T^79^ using BWA-mem2 (Version: 2.2. 1) with default parameters. Mapping results were converted into the BAM format and filtered for unmapped and non-unique reads using SAMtools (Version: 1.17). BWA alignment was conducted, and two functions “vc and joint” of GTX (Version: 2.2. 1) were applied to obtain SNP VCF files. To ensure the accuracy of the results, SNPs with a missing genotype frequency greater than 0.05 or a minor allele frequency (MAF) less than 0.05 were excluded from analysis. Imputation was not performed. Populations were considered for their structure and cryptic relationships, and GEMMA (Version: 0.98. 3) was used to implement the GWAS of the standard linear mixed model:

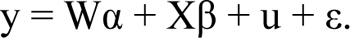

The whole-genome significance cutoff was defined using the adjusted Bonferroni test threshold, which was set as *P* < 1e-5.

### Transcriptomic analyses under herbivorous conditions

Our transcriptome data sets are originated from the Sequence Read Archive (SRA) accession SRP067967, which was sequenced for its transcriptome data on grape leaves under herbivorous exposure to two-spotted spider mite (*Tetranyhus urticae*)^39^. The raw transcriptomic data was processed using fastq-dump (Version:2.11. 0), converting it into paired-end fastq files. Quality control was performed using FastQC (Version:0.11. 9) to assess data quality. Subsequently, trim-galore software (Version:0.6. 7) was utilized to filter out adapter sequences and low-quality reads (--Stringency 3--phred33 -q 25). The data was then aligned to the grape reference genome using STAR software (Version:2.7. 10b), resulting in aligned BAM files. Quantification of gene expression was performed using featureCounts (Version:2.0. 1). For the final analysis, we employed DESeq2 package (Version:1.38. 3) in RStudio to conduct differential gene expression analysis. Genes with a fold change greater than two and a *P* value less than 0.05 will be considered as differentially expressed genes (DEGs). All GO enrichment analyses were conducted on the DAVID website (https://david.ncifcrf.gov/tools.jsp). Additionally, we employed materials from BioRender (https://app.biorender.com/) to construct mode pattern.

### The comparison of genomic selection models based on the Cross-validation

Regarding the severity of pest damage, two phenotypic descriptors were estimated. One was binary qualitative shape with the phenotype divided into two classes (“mild” vs. “severe”). The other one represented a quantitative measurement of the damage with the phenotype quantified into a continuous scale ranging from 0 to 5. For binary phenotype, Logistic Regression, Support Vector Classifier (SVC), Random Forest Classifier and Naive Bayes Classifier models were selected to enable comparisons of predictive accuracy and bias. The prediction accuracy between different models was determined by the average accuracy. For continuous phenotype prediction, two machine learning models, Support Vector Regressor (SVR), Random Forest Regressor (RFR), and three regularization models, Lasso (L1 regularization), Ridge regression (L2 regularization) and ElasticNet (combining L1 and L2 regularization), were compared between each other. L1 and L2 regularization represent two types of marker effect estimation. The former, L1 regularization, is well-suited for identifying and adapting to a few major QTLs that have significant effects on the phenotype, L2 regularization, also known as ridge regression, is effective in capturing the effects of many minor-effect QTLs. The Pearson’s correlation coefficient between the predicted and the observed phenotypes were calculated as the prediction accuracy. The support vector machine (SVM) models were applied to predict both binary and continuous phenotypes. Additionally, SVM offers various kernel functions, including the linear kernel, polynomial kernel, and radial basis function kernel. These three kernel functions were also compared by the cross-validation to select the optimal one that represented the best results for this model. The all models utilized in genomic selection were implemented using the scikit-learn Python library (Version: 1.0. 2). Different models in two types of pest phenotypes were compared based on accuracy and the optimal model was chosen.

### Stepwise variation selection based on GWAS results for optimal prediction

Regarding the results from GWAS, the sorting based on their *P* values and a stepwise selection of variations were performed to identify the optimal number of variations to construct a genomic selection model. Before modeling, Plink (Version:1.90b7) was used to filter variations with the parameters including window size set to 100, step size set to 50, and linkage disequilibrium (LD) threshold set to 0.2 to reduce redundancy and minimize the impact of LD in the genomic selection process. After filtering, a total of 7,359,942 variations were reduced to 313,961 variations, representing all the variations participated in the modeling. Different numbers of variations in models with binary and continuous phenotypes were evaluated with cross validation. With the binary phenotype, a total of 50 significant SNPs (*P* < 1e-5) were identified through GWAS. The variations with the numbers including 1, 10, 50 (all significant SNPs), 100, 500, 2000, 5000, and 313,961 (all variations) according to their sorted *P* values were selected for genomic selection. Regarding the continuous phenotypes, a total of 53 significant SNPs (*P* < 1e-5) were identified. The numbers of variations including 10, 53 (all significant SNPs), 100, 500, 2000, 5000, 10000, and 313,961 (all variations) were considered for prediction modeling. 10% of the samples’ phenotype data were reserved for testing, while the remaining 90% of the samples were split in a 4:1 ratio for modeling and validation. The optimal number of variations was evaluated based on the final modelling accuracy averaged over one hundred iterations. Random seeds were set to ensure the training data consistency across these 100 modeling and evaluation iterations.

## Data availability

The raw data in this study have been deposited into in zenodo (XXX).

## Code availability

All scripts performed in this study are available on Github: https://github.com/zhouyflab/Pest-Resistance

## Acknowledgements

We thank members of the Zhou lab at AGIS for discussion and comments on the project. This work was supported by the National Key Research and Development Program of China (2023YFF1000100) and the National Natural Science Fund for Excellent Young Scientists Fund Program (Overseas) to Yongfeng Zhou.

## Author contributions

Y. Z. designed the project. Y. Z., Z. L. and Y. W. supervised the project. Y. G., Z. L. design and train image recognition and object detection models. Y. G., F. Z., and Q. X. performed the bioinformatic analyses. X. W., H. X., W. L., L. S., Y. Z., and C. L. assisted in bioinformatics analyses., Y. G., X. S., and W. M. provided essential technology support. Q. L. and A. M. provided the plant material for genome analyses. Y. G., Z. L., and Y. W. wrote the draft. Y. L., X. F., X. F., C. Z., P. Z., C. L., and Z. Z. revised the manuscript. All authors contributed to manuscript preparation and read, commented on, and approved the manuscript.

