## Supplementary Fig 1-12 for "Deep learning based genomic breeding of pest-resistant grapevine"

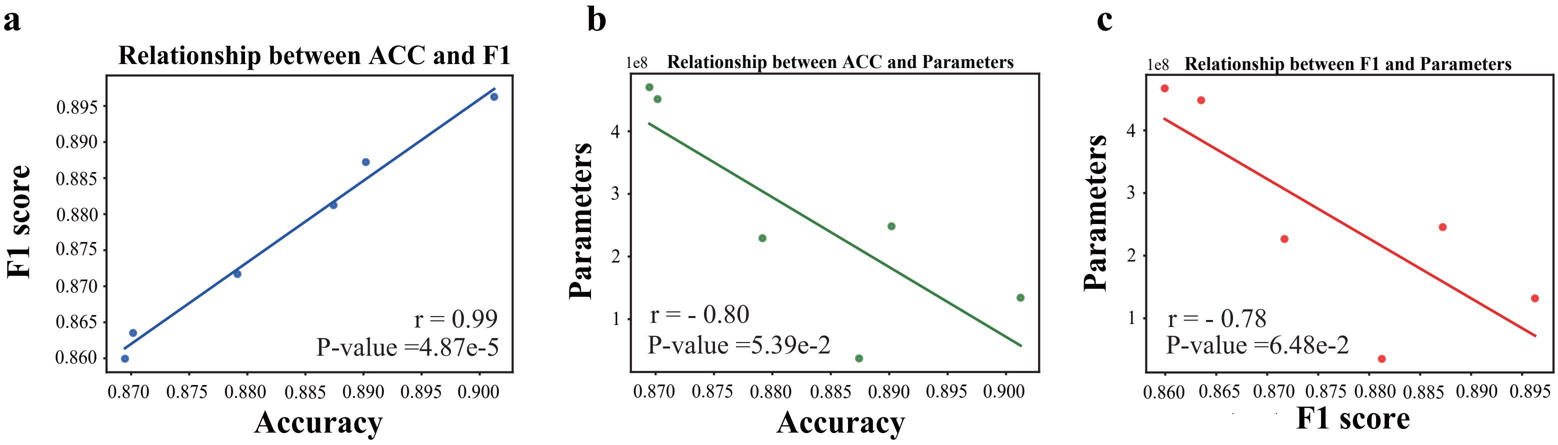


**Supplementary Fig. 1 | Analysis of the relationship between F1 score, accuracy and paramters. a**, A scatter plot shows the average accuracy and F1 scores obtained from six different models in cross-validation,along with thier correlations. **b**, Correlations and a scatter plot illustrating the accuracy obtained from six models in relation to the number of training parameters. **c**, Correlations and a scatter plot illustrating the F1 score obtained from six models in relation to the number of training parameters.

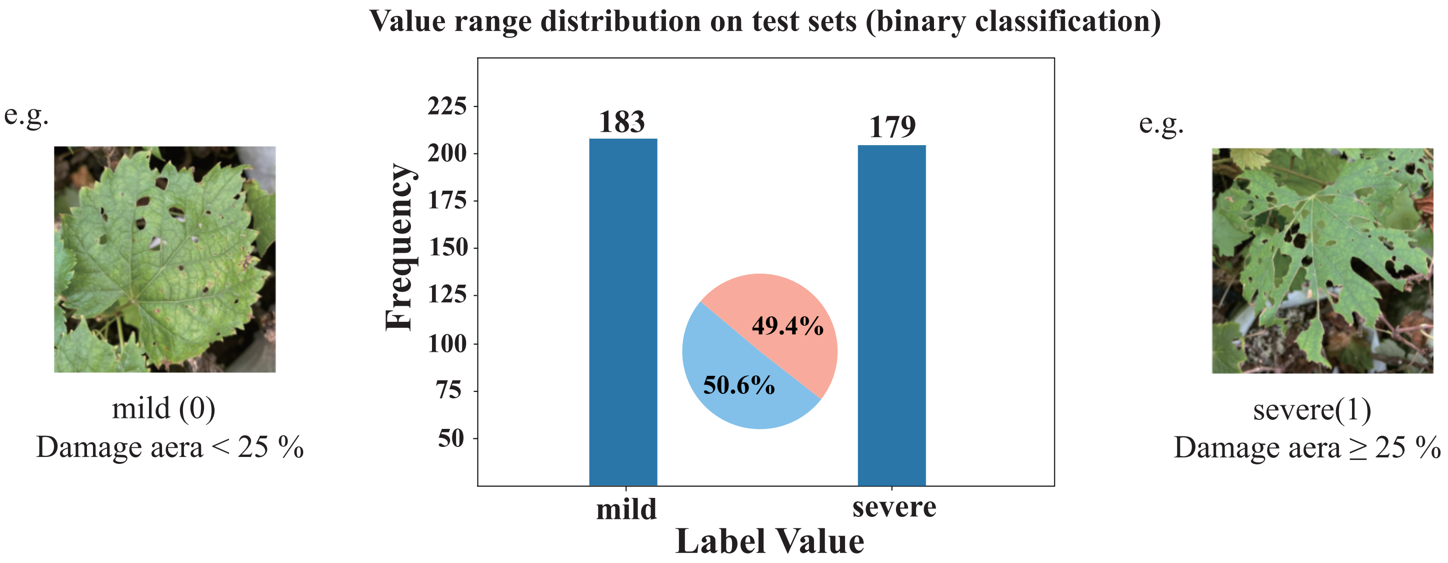


**Supplementary Fig. 2 | Numerical distribution of test datasets for classification models.** The numerical distribution of the test set used by VGG16, The examples of severe and mild pest damage on leaves are shown on the left and right, respectively. The bar chart and pie chart depict the specific quantities and proportions of different categories, illustrating the severity levels of the pest infestation.

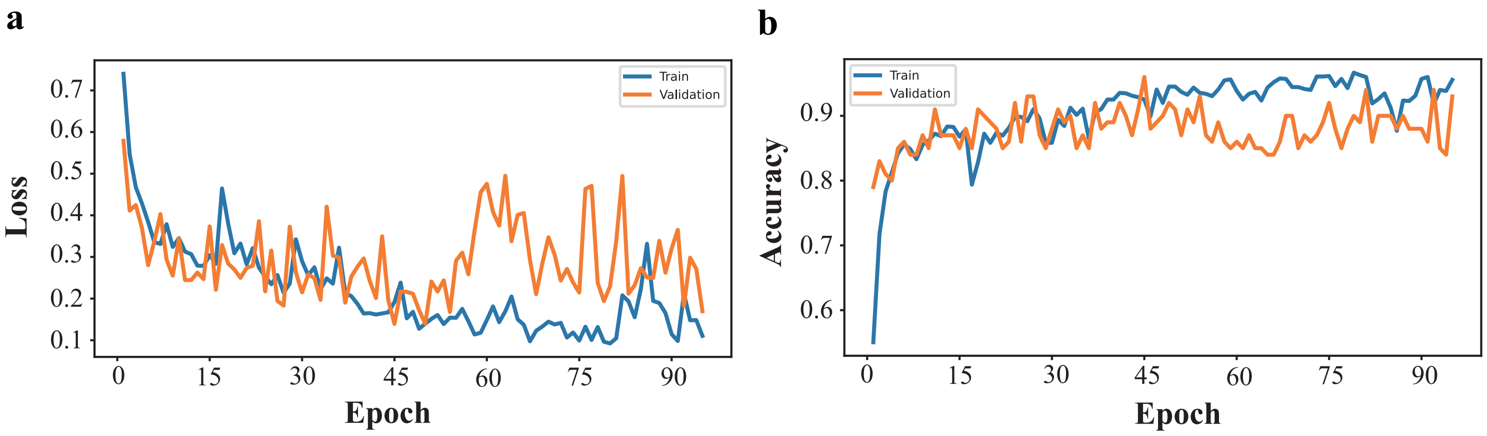


**Supplementary Fig. 3 | Learning curve of VGG16. a**, The fluctuations in the loss function during VGG16 training. **b**, The changes in accuracy during VGG16 training.

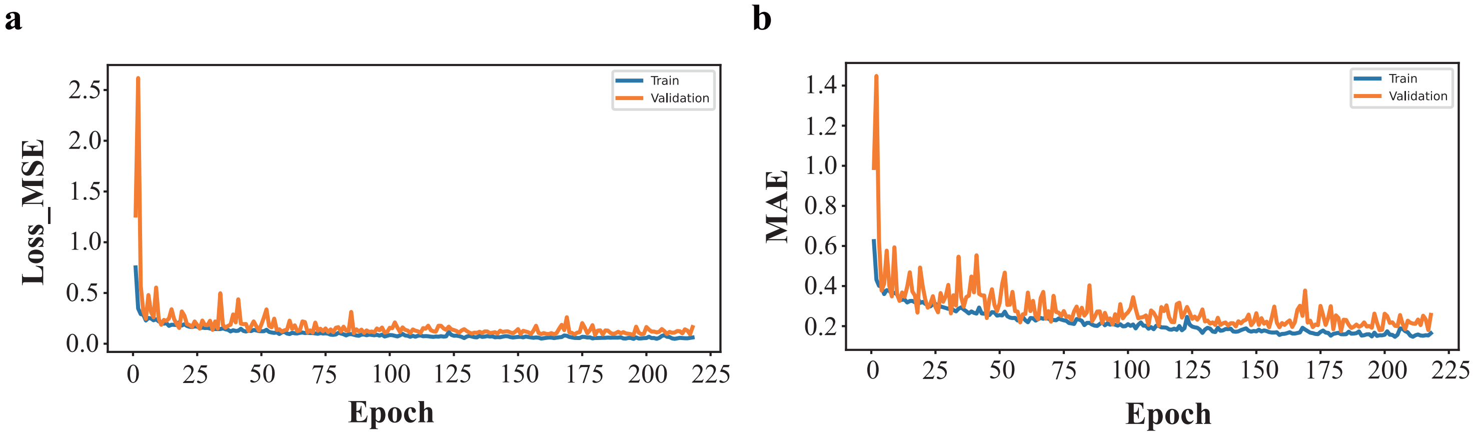


**Supplementary Fig. 4 | Learning curve of DCNN-PDS. a**, Changes in loss function (MSE) during VGG16 training. **b**, Changes in MAE during DCNN-PDS training.

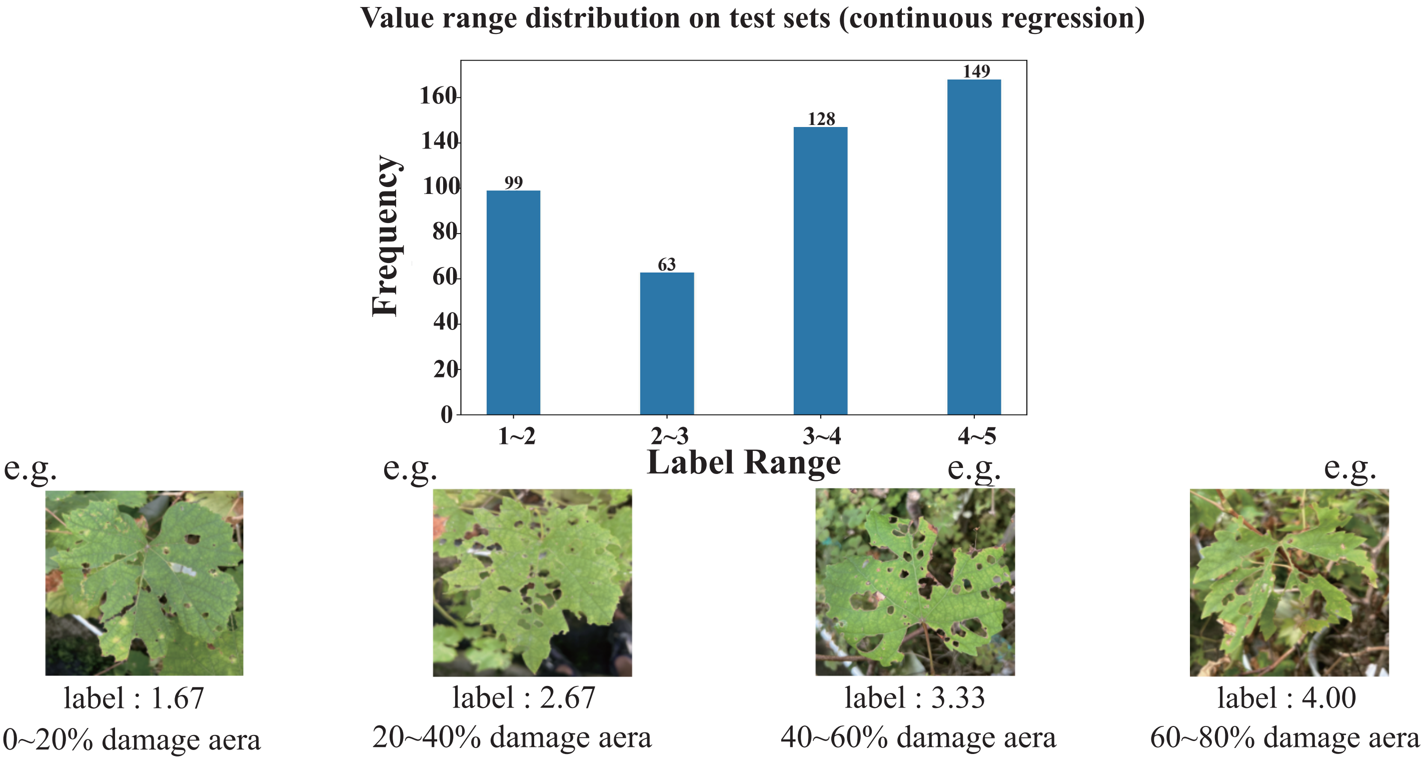


**Supplementary Fig. 5 | The numerical distribution of the test set used by DCNN-PDS.** The bar chart illustrate the quantity of leaf images labeled for various pest severity levels by different raters, while the bottom section displays some sample examples.

**a**

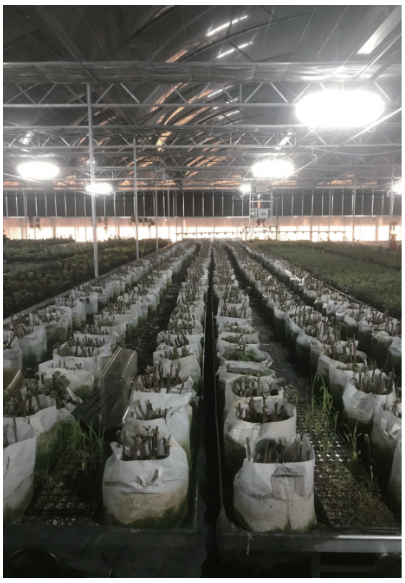

**b**

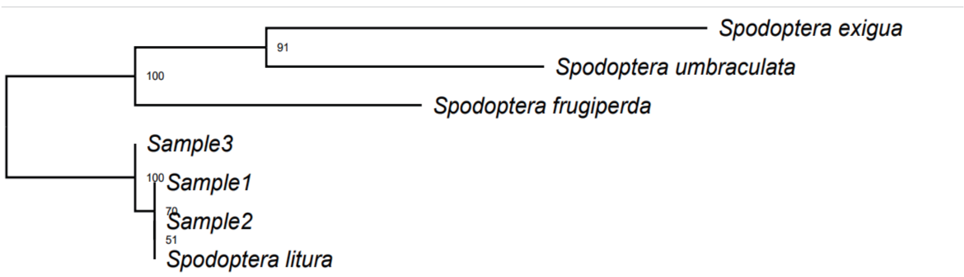


**Supplementary Fig. 6 | Layout and enviromental conditions of greenhouse viticulture and experimental pest situation. a**, Plant and cultivation of different grape varieties under greeenhouse conditions. **b**, The evolutionary tree constructed for experimental pests is situated within the same branch as the Spodoptera litura.

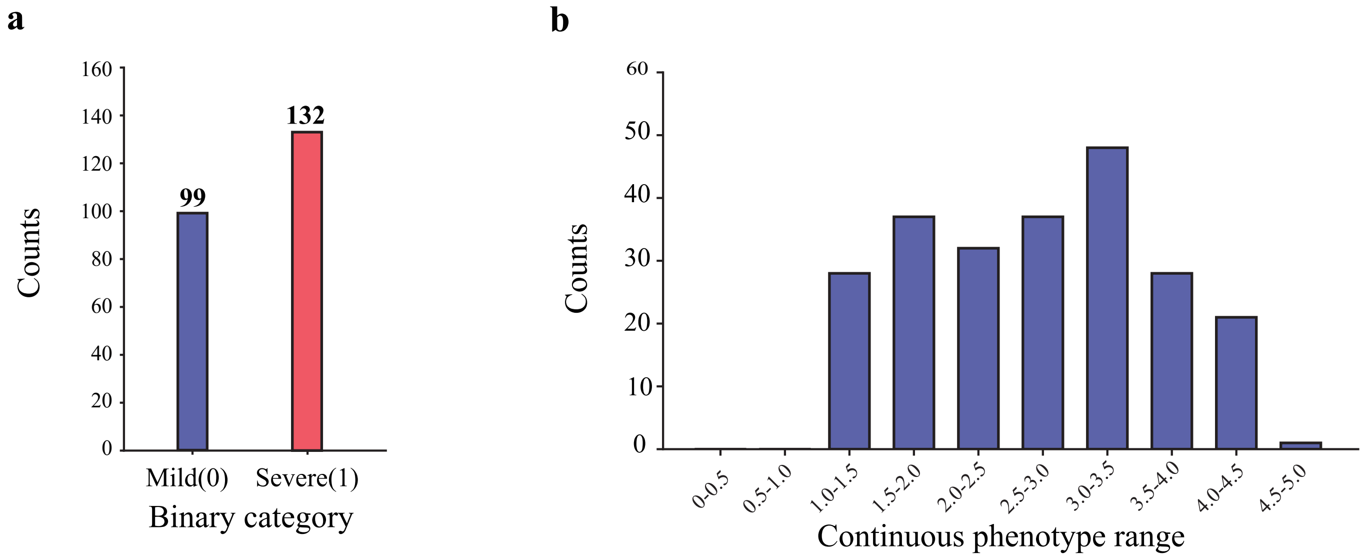


**Supplementary Fig. 7 | The overall distribution of binary and continuous phenotypes among 231 varieties. a**, The binary phenotypic status of herbivore damage measured across all tested accessions. **b**, The overall distribution of continuous phenotypes.

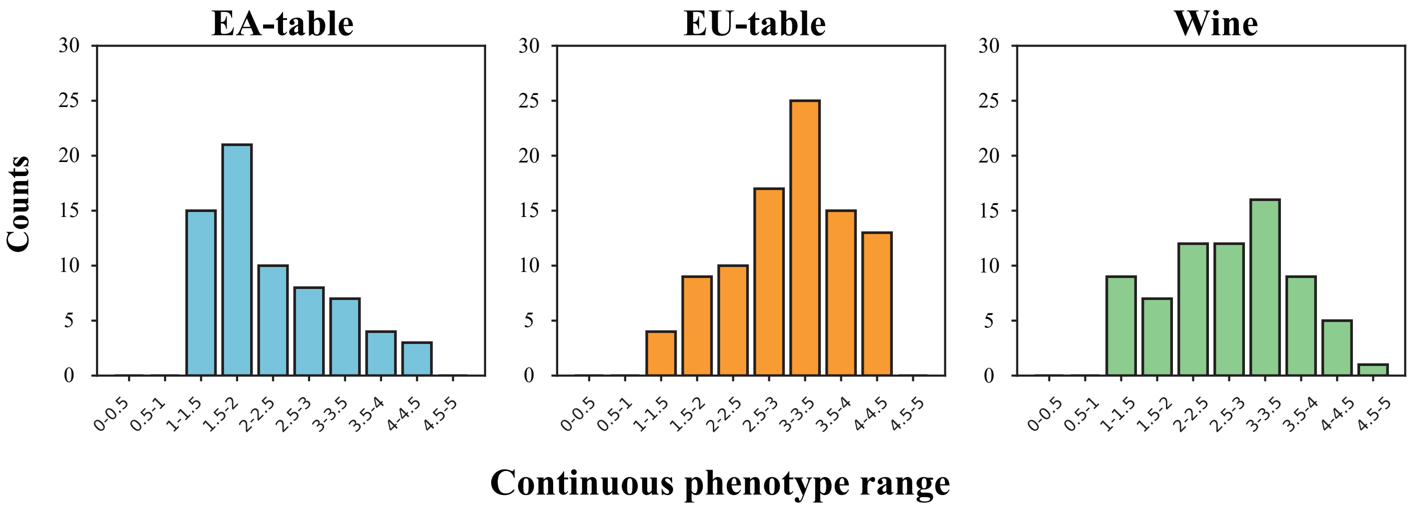


**Supplementary Fig. 8 | The overall distribution of continuous phenotypes in three different categories of grapes.** From left to right, it represents the distribution of continuous phenotypes in three different categories of grapes: Euro-American table grapes, Euro-Asian table grapes, and Euro-Asian wine grapes.

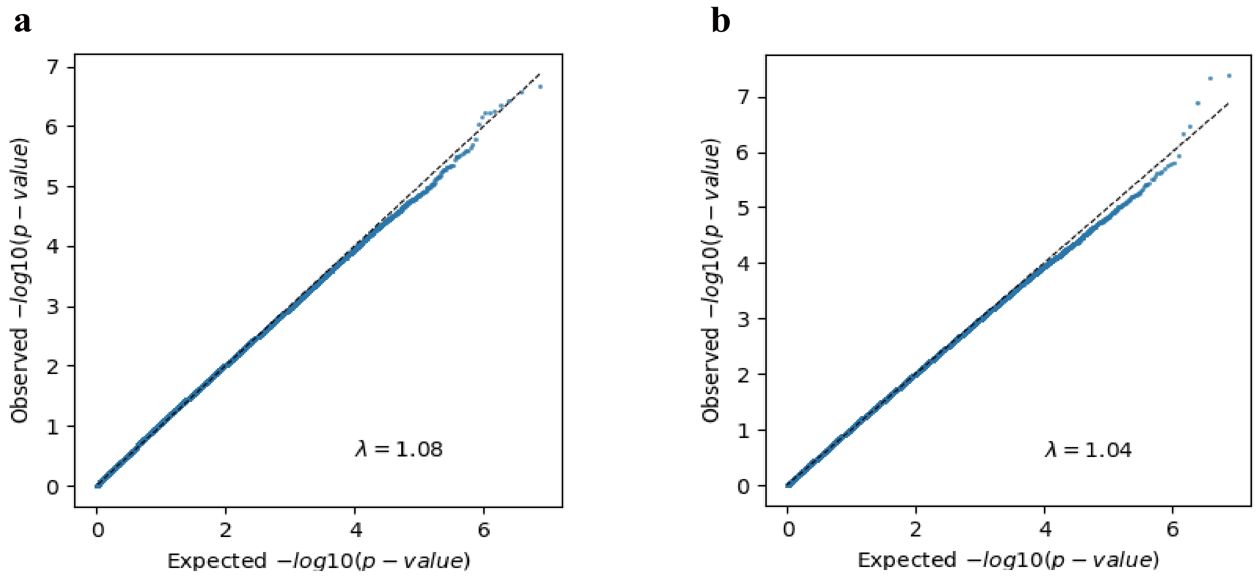


**Supplementary Fig. 9 | QQ plot of GWAS results. a**, Binary traits of GWAS result. **b**, Continuous traits of GWAS result.

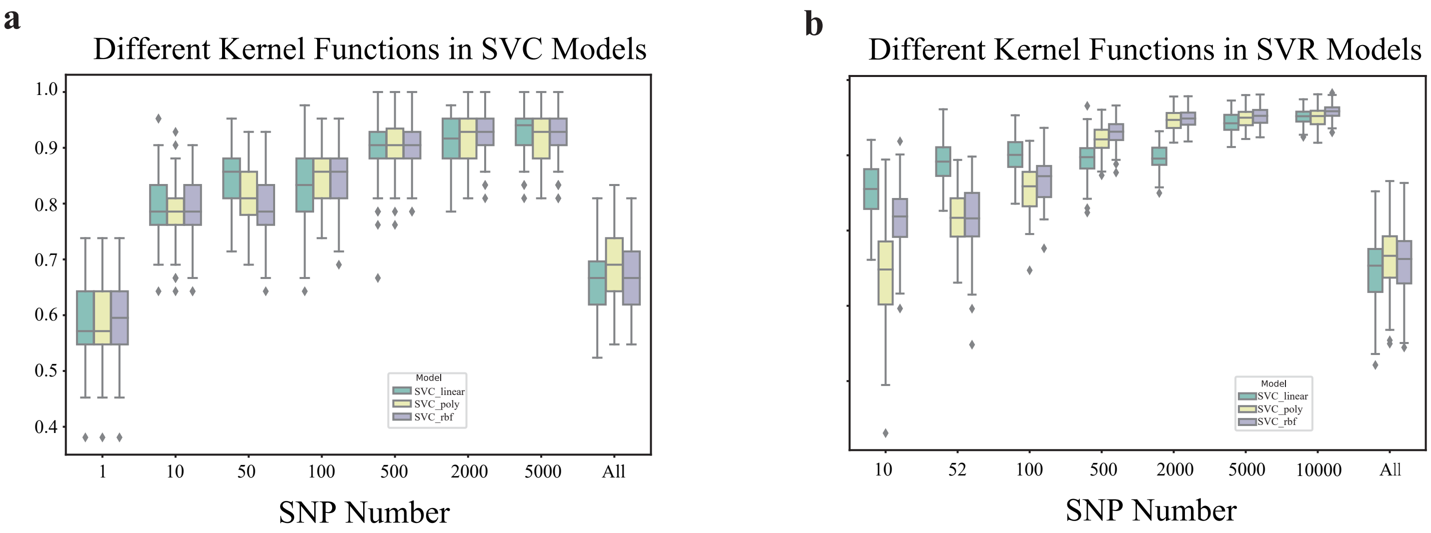


**Supplementary Fig. 10 | Cross-Validation of Support Vector Machine Models with Different Kernel Functions. a**, Performance of support vector classification (SVC) models with different kernel functions in predicting binary phenotypes. **b**, Performance of support vector regression (SVR) models with different kernel functions in predicting continuous phenotypes.

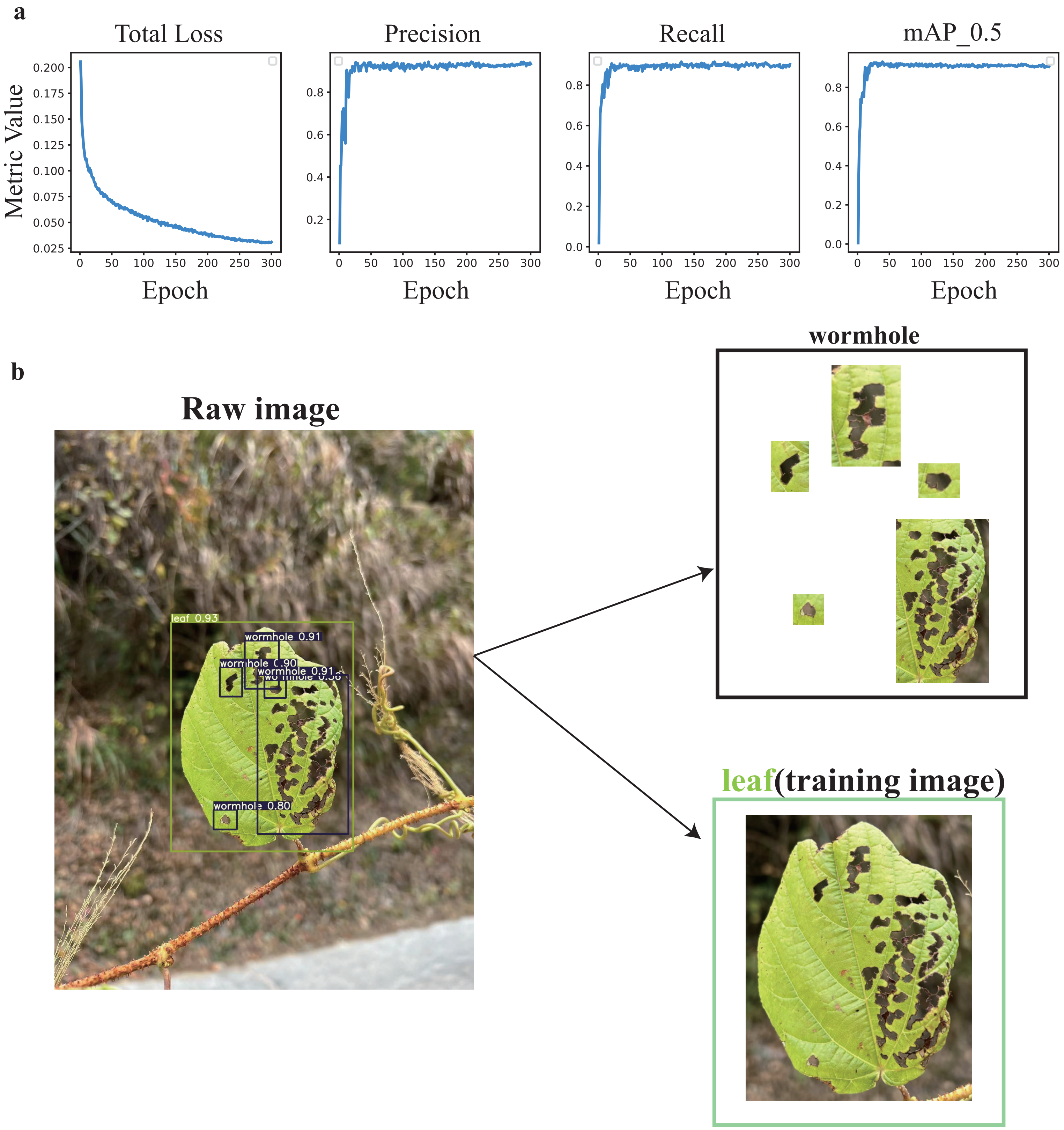


**Supplementary Fig. 11. YOLOv5 Object Detection Training Curves and Image Processing. a**, the YOLO model shows the change curves for total loss, precision, recall, and mAP_0.5 in 300 epochs of training. **b**, After performing object detection on the original images, leaf target boxes are extracted as the final training images

| Sample | 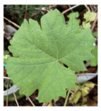 | | 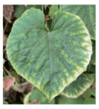 | | 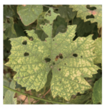 | | 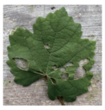 | | 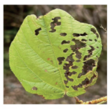 | | 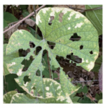 | | 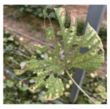 | | 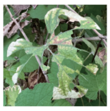 | | 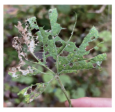 | | 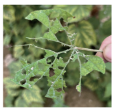 | |
|---|---|---|---|---|---|---|---|---|---|---|---|---|---|---|---|---|---|---|---|---|
| Rater1 | 0 | 1 | 0 | 1 | 0 | 1 | 0 | 1 | 1 | 2 | 1 | 2 | 1 | 3 | 1 | 4 | 1 | 4 | 1 | 5 |
| Rater2 | 0 | 1 | 0 | 1 | 0 | 1 | 1 | 2 | 1 | 2 | 1 | 3 | 1 | 4 | 1 | 4 | 1 | 5 | 1 | 5 |
| Rater3 | 0 | 1 | 0 | 1 | 0 | 2 | 0 | 2 | 0 | 3 | 1 | 3 | 1 | 4 | 1 | 5 | 1 | 5 | 1 | 5 |
| Binary_label | 0 | | 0 | | 0 | | 0 | | 1 | | 1 | | 1 | | 1 | | 1 | | 1 | |
| PDS_label | 1 | | 1 | | 1.33 | | 1.67 | | 2.33 | | 2.67 | | 3.67 | | 4.33 | | 4.67 | | 5 | |

**Supplementary Fig. 12 | Binary and Continuous Labeling Methods and Examples.** Each image received two ratings from the raters a binary rating (left) and a continuous level rating (right). The binary label are determined by Majortiy Vote, where if two raters provide the same rating, their ratings are adopted as the label values. The continuous label (PDS_label) are determined by the average of the scores from three raters.
